# Macroevolutionary analysis of discrete character evolution using parsimony-informed likelihood

**DOI:** 10.1101/2020.01.07.897603

**Authors:** Michael C. Grundler, Daniel L. Rabosky

**Affiliations:** Museum of Zoology and Department of Ecology and Evolutionary Biology, University of Michigan, Ann Arbor, MI 48109-1079, USA

**Keywords:** macroevolution, rate variation, parsimony, likelihood, character evolution

## Abstract

Rates of character evolution in macroevolutionary datasets are typically estimated by maximizing the likelihood function of a continuous-time Markov chain (CTMC) model of character evolution over all possible histories of character state change, a technique known as maximum average likelihood. An alternative approach is to estimate ancestral character states independently of rates using parsimony and to then condition likelihood-based estimates of transition rates on the resulting ancestor-descendant reconstructions. We use maximum parsimony reconstructions of possible pathways of evolution to implement this alternative approach for single-character datasets simulated on empirical phylogenies using a two-state CTMC. We find that transition rates estimated using parsimonious ancestor-descendant reconstructions have lower mean squared error than transition rates estimated by maximum average likelihood. Although we use a binary state character for exposition, the approach remains valid for an arbitrary number of states. Finally, we show how this method can be used to rapidly and easily detect phylogenetic variation in tempo and mode of character evolution with two empirical examples from squamates. These results highlight the mutually informative roles of parsimony and likelihood when testing hypotheses of character evolution in macroevolution.

## Introduction

Continuous-time Markov chains (CTMC) are commonly used in macroevolution to model the evolution of discrete characters to make inferences about evolutionary rates and patterns of change. Because the historical sequence of ancestor-descendant character state changes in most cases is not directly observed, statistical inference about Markov chain models is typically performed by summing over all possible histories of character evolution that could have resulted in the observed distribution of character states among living species using the well-known peeling algorithm (Felsenstein 1981). This technique, sometimes referred to as “maximum average likelihood” (Steel & Penny 2000), weights each possible evolutionary pathway by its probability of generating the observed data, and the resulting parameter estimates are therefore not conditioned on any particular history of character evolution. Pagel (1999) terms these “global estimators” and they are recommended as best practice by Mooers & Schluter (1999) and the only implementation available in most commonly used software packages in macroevolution (e.g., Paradis et al. 2004; Revell 2012; FitzJohn 2014).

An alternative approach to maximum average likelihood, termed “most-parsimonious likelihood”, is to estimate both the unknown internal node states and the unknown transition rates simultaneously using likelihood (Barry and Hartigan 1987). Goldman (1990) refers to internal node states as “incidental parameters”, since they are realizations of a random process, and raises concerns about the statistical consistency of such estimates. Perhaps as a result, most-parsimonious likelihood techniques do not appear to be used in common practice. Alternatively, internal node states can be estimated independently (using parsimony), and likelihood estimates of transition rate parameters can be made by conditioning on these possible parsimonious pathways of evolution. In a macroevolutionary context, Janson (1992) appears to have been the first to use such an approach in a study of seed dispersal syndromes in plants. Some properties of these estimators were later studied by Sanderson (1993). The approach has received little attention since these two early works.

Here we revisit the approach of conditioning transition rate estimates on parsimonious ancestral state reconstructions, an approach we term parsimony-informed likelihood. We find that transition rates estimated using parsimony-informed likelihood have lower mean-squared error than rates estimated using maximum average likelihood and that this difference is accentuated in simulations where likelihood-ratio tests support an asymmetric model over a symmetric model of character evolution. We then show how the simple form of the parsimony-informed likelihood estimator leads to a rapid way of exploring datasets for phylogenetic variation in tempo and mode of character evolution.

## Materials & Methods

### General theoretical background

Given an assignment of character states to each node in a phylogeny the likelihood of the observed character state data at the terminal nodes is the product of state-to-state transition probabilities over all ancestor-descendant pairs. The full likelihood is computed by summing these evolutionary pathway likelihoods over all possible histories of character evolution that could have given rise to the observed character state data using Felsenstein’s (1981) peeling algorithm. We refer to transition rates estimated with this method as maximum average likelihood rates, abbreviated MAV. In the general multistate case, the transition probabilities used in the peeling algorithm are computed by solving the matrix exponential exp(***Q****t*), where ***Q*** is the matrix of transition rates and *t* is the branch length separating ancestor and descendant.

However, the simple two-state case like the kind used in this study permits an analytic solution given by,

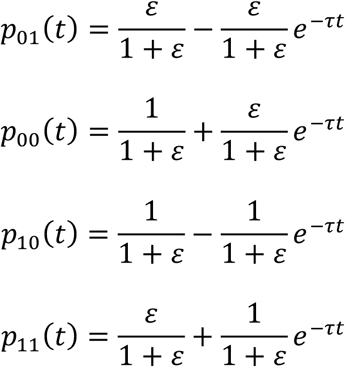

where 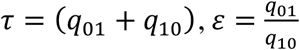, and *q*_01_ and *q*_10_ are the forward and reverse transition rates, respectively.

Parsimony-informed likelihood begins by restricting attention only to the histories of character of evolution inferred with maximum parsimony (MP). There will often be more than one MP history and the set of MP histories will be a subset of the character histories used in the computation of the maximum average likelihood. By treating these ancestor-descendant state reconstructions as “observations” sampled from a CTMC at random time points we can estimate the transition probability matrix of the discrete-time Markov chain embedded within the CTMC. This is,

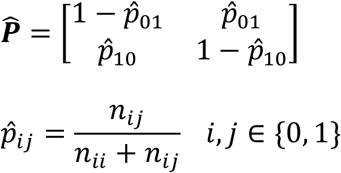

Here, 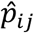 is the probability of a gain or loss and is calculated as the number of branches where a change from *i* → *j* occurred divided by the number of branches where such a change could have occurred. In practice, these probabilities are calculated by sampling MP histories of character evolution using Fitch’s three algorithms (1971). We estimated this matrix by sampling 1000 MP histories such that each permissible state-to-state transition had uniform weight. In actuality, the number of MP histories can be much larger than this. If we further assume that the random time points at which we “observe” the CTMC are independent of the process of character evolution and distributed according to a Poisson process, we can use 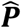 to estimate the transition rate matrix ***Q*** of the CTMC. This is,

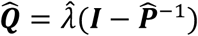

Where 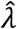 is the estimated rate of the Poisson process that generates the time points at which we observe the chain’s state. In this study, we take 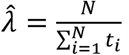, the number of nodes divided by the summed branch length of the phylogeny. For a two-state model 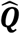 is simply,

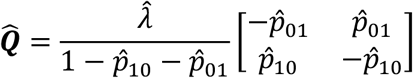

This estimator is derived by Karr (1991, pp. 384-386). It was used by Janson (1992) in a study of plant dispersal syndromes that remains a lucid but under cited discussion of the role of Markov models in macroevolutionary inference. Although we present results for only two states the method is applicable to any number of states (i.e., for *K* states 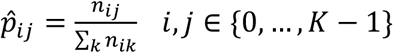).

Note that 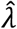 does not influence the relative values of the estimated rate matrix. Only the pattern of transitions (including apparent stasis) affects the relative values. This is an important difference from the MAV estimator used in standard practice, which is also affected by variation in branch lengths over which transitions occur. We refer to rates estimated with this method as parsimony-informed likelihood transition rates, abbreviated PIL.

From a computational perspective PIL has a clear advantage over MAV because it requires only a single matrix inversion instead of repeated matrix exponentiation, which makes PIL more feasible for datasets with many character states. A disadvantage of PIL is that it can lead to undefined transition rates when change is rare enough that not all possible transitions are observed in parsimony reconstructions. A second potential advantage is that 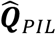 may be less variable and therefore less prone to error than 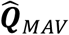 because 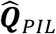 is conditioned on a minimal set of character state changes needed to explain the data. We explore this possibility in the simulations described below.

### Simulations

We simulated 100 single-character datasets on each of 50 empirical phylogenies using a two-state CTMC model of discrete character evolution. To ensure simulated character datasets were evolved on phylogenies with realistic branch length distributions, we obtained empirical phylogenies by randomly sampling clades containing between 100 and 1,000 terminal nodes from a recent phylogeny of ray-finned fishes (Rabosky et al. 2018). For each simulation we chose model parameters *ε* and *τ* such that the probability of state identity between every ancestor and descendant pair was at least one-half. In other words, simulations were parameterized such that a change of state from ancestor to descendant was never more probable than retaining the ancestral state. This property does not preclude simulations from having a fast rate of evolution, but it does require that transition rates become more symmetrical as the rate of evolution increases. For each simulated dataset we estimated *ε* and *τ* using the MAV and PIL estimators discussed above. When performing MAV estimates we used both an unconstrained and a constrained (*ε* = 1) model. Our rationale for fitting a constrained model to simulated datasets is that in practice researchers will often select an unconstrained model for analysis only when a likelihood ratio test indicates it substantially improves model fit over a constrained model. We conducted all simulations and parameter estimation in R 3.5.2 (R Core Team 2018) using the package macroevolution (available from https://github.com/blueraleigh/macroevolution).

### Variation in tempo and mode

Because 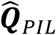 can be computed relatively rapidly it presents a simple method for detecting phylogenetic variation in tempo and mode of character evolution. First, compute 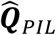 for the entire tree and for each clade showing some minimum number of parsimony-inferred character state changes. If the minimum number of parsimony changes required for the test is too low rates cannot be estimated, but if it is too high there will be no opportunity to test for variation in tempo and mode. We used a value of 10 in our analysis. Second, for each of these clades compute the maximum average likelihood of their data using their clade-specific 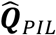 as well as the tree-wide 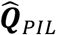. The magnitude of the log-ratio formed from these two likelihoods will be an indicator of how strongly the rate and pattern of character state changes in the clade differs from the rest of the phylogeny. For a two-state character the log-ratio should be at least 4 before concluding that a clade displays a significantly different tempo and mode of character evolution than the rest of the phylogeny. This is only a heuristic since we do not actually fit a model with multiple rate matrices. Its value can be derived by assuming the clade-specific log-likelihood ratio represents the difference in log-likelihoods between a model with and without a clade-specific rate-matrix and by requiring a minimum difference of zero in the AIC between these two models before accepting the model with the clade-specific rate-matrix.

## Results

Across all simulations the mean squared error (MSE) in 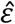 for MAV (MSE = 1.86) and PIL (MSE = 1.82) estimators was very similar (Fig. 1). However, comparing just the subset of simulations for which an unconstrained model outperforms a constrained model by a likelihood-ratio test reveals dramatic differences, with MSE for PIL estimates being substantially lower (MSE = 1.82) than for MAV estimates (MSE = 6.99) (Fig. 1). The cause of this difference is evidently from the tendency of the likelihood-ratio test to select a subset of simulations that show clearly biased MAV estimates. When comparing 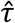 across all simulations, unconstrained MAV (MSE = 1.38) and constrained MAV (MSE = 1.18) estimates had higher MSE than PIL (MSE = 0.0012) estimates (Fig. 2). These differences also persisted in the subset of simulations for which an unconstrained model outperforms a constrained model by a likelihood-ratio test, with unconstrained MAV estimates (MSE = 2.80) having the highest MSE, followed by constrained MAV estimates (MSE = 0.35), and with PIL estimates (MSE = 0.00081) again achieving the lowest MSE.

**Figure 1.**
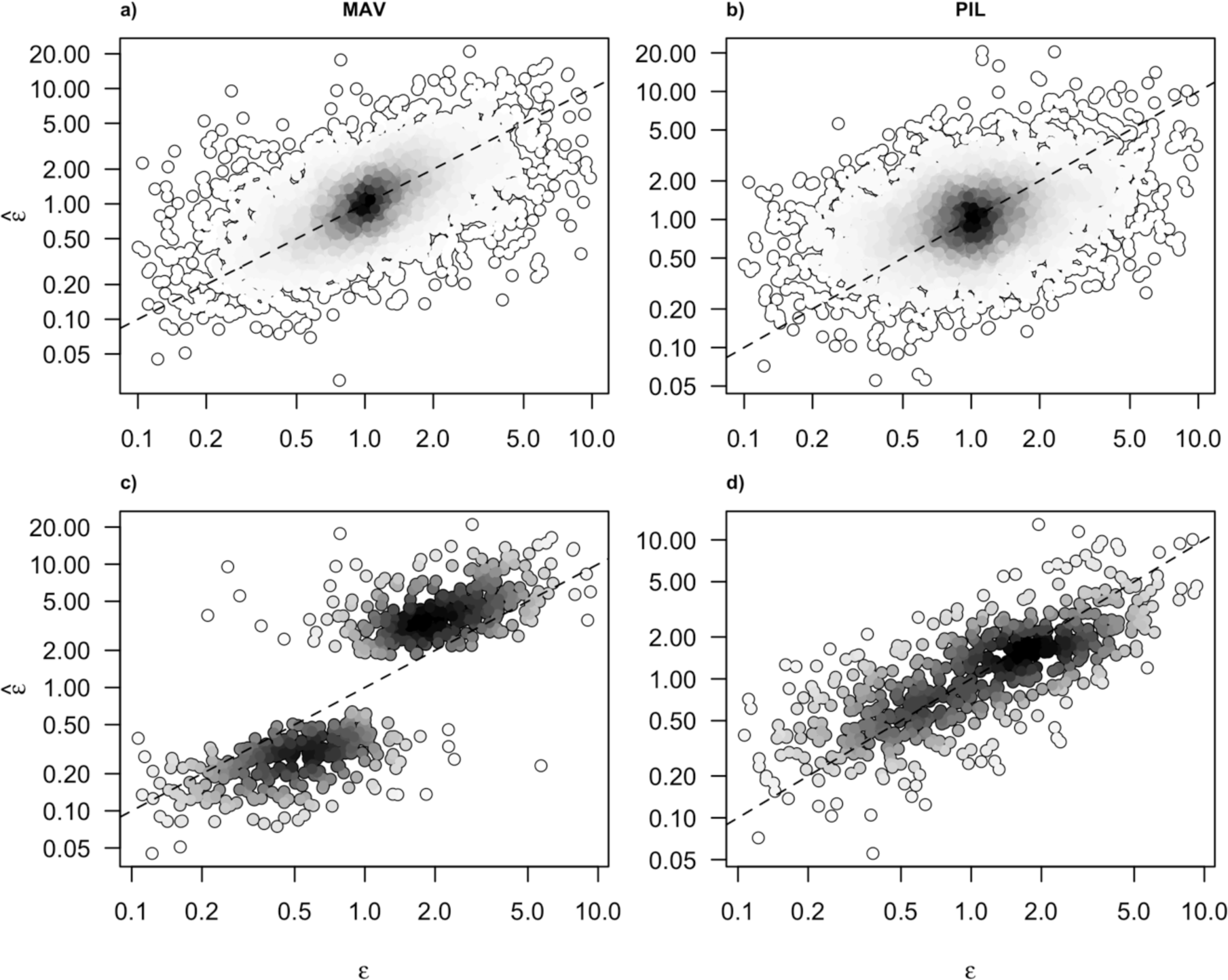
Relationship between the true (*ε*) and estimated 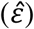 ratio of forward and reverse transition rates for maximum average likelihood (MAV) and parsimony-informed likelihood (PIL) estimators. Darker shading indicates a higher density of points. The top panels show estimates for the full set of simulations. The bottom panels show estimates for the subset of simulations for which a likelihood ratio test determined an unconstrained model offered a significantly better fit than a constrained (*ε* = 1) model.

**Figure 2.**
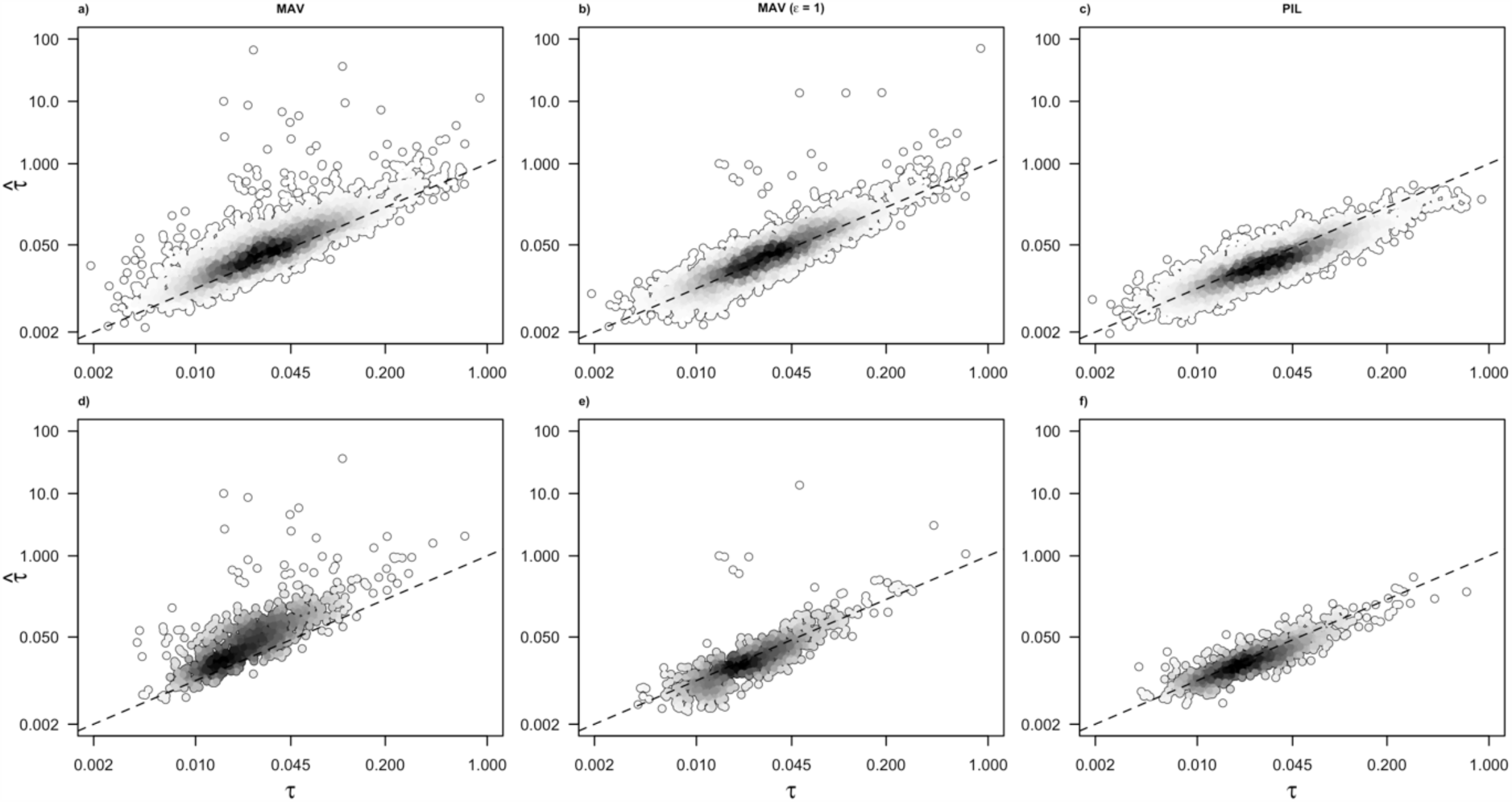
Relationship between the true (*τ*) and estimated 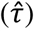 sum of forward and reverse transition rates for maximum average likelihood (MAV) and parsimony-informed likelihood (PIL) estimators. Darker shading indicates a higher density of points. The top panels show estimates for the full set of simulations. The bottom panels show estimates for the subset of simulations for which a likelihood ratio test determined an unconstrained model offered a significantly better fit than a constrained (*ε* = 1) model.

When we used PIL to detect phylogenetic variation in tempo and mode in datasets simulated with no variation in tempo and mode, from 0% – 11% of simulations showed clades with significant departures in tempo and mode from the rest of the phylogeny. Pooling simulations across all phylogenies, only 2.6% of simulations detected significant phylogenetic variation in tempo and mode. Using stricter criteria (i.e. requiring more character state changes and a higher log-likelihood ratio) would further lower these percentages. These values indicate that the test does not readily identify variation in tempo and mode where none exists. When we applied the test to two previously analyzed empirical datasets it identified sets of clades showing variable tempo and mode that were in broad agreement with previous analyses that used computationally intensive MCMC techniques for the same purpose (Fig. 3, 4). We note that each of these two empirical applications of the test requires only a few seconds, permitting rapid exploration of datasets for clades with variable tempo and mode.

**Figure 3.**
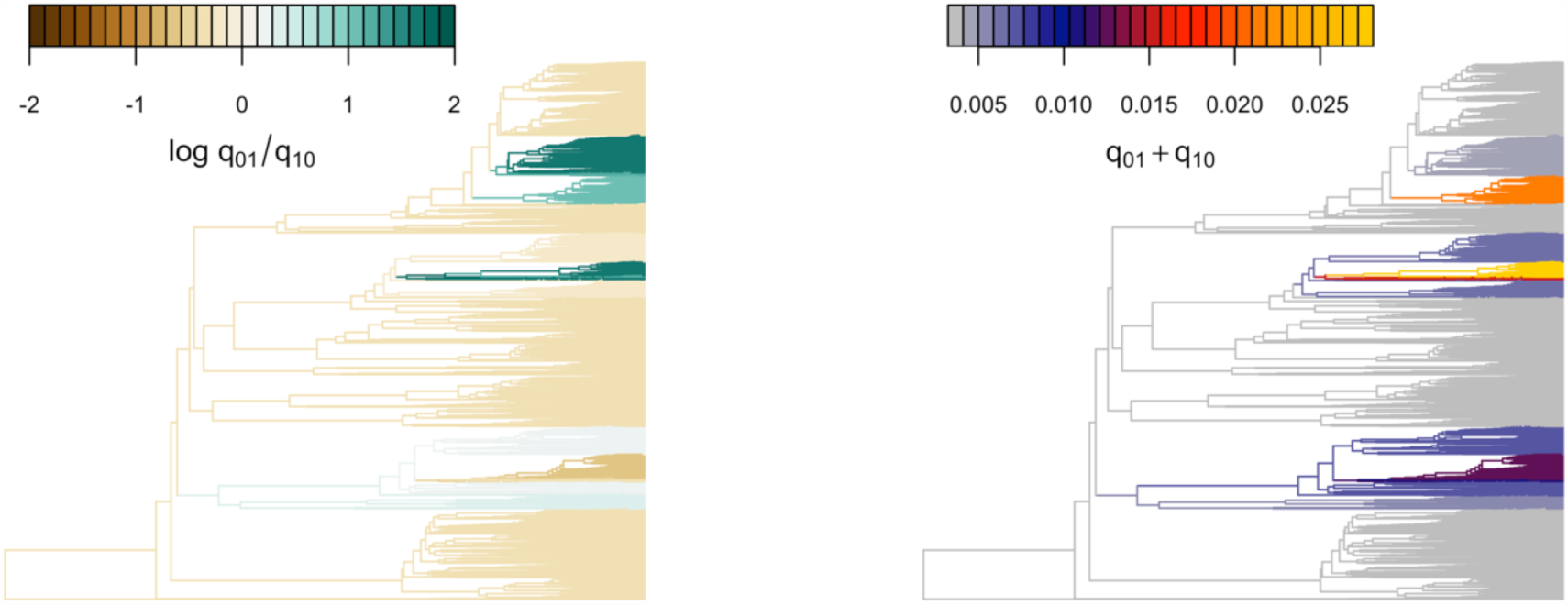
Evolutionary gains and losses of viviparity show variable tempo and mode across the squamate tree of life. Rate matrices were estimated using maximum likelihood conditioned on histories of character evolution inferred with parsimony for clades with at least 10 parsimony-inferred changes. Each clade-specific rate matrix was then compared to the rate-matrix for the whole tree. If the log-likelihood of the data in the subtree formed by a clade was at least 4 log-likelihood units greater under its clade-specific rate matrix than under the tree-wide rate matrix its branches were colored according to the parameters of that matrix. Data from Pyron and Burbrink (2014).

**Figure 4.**
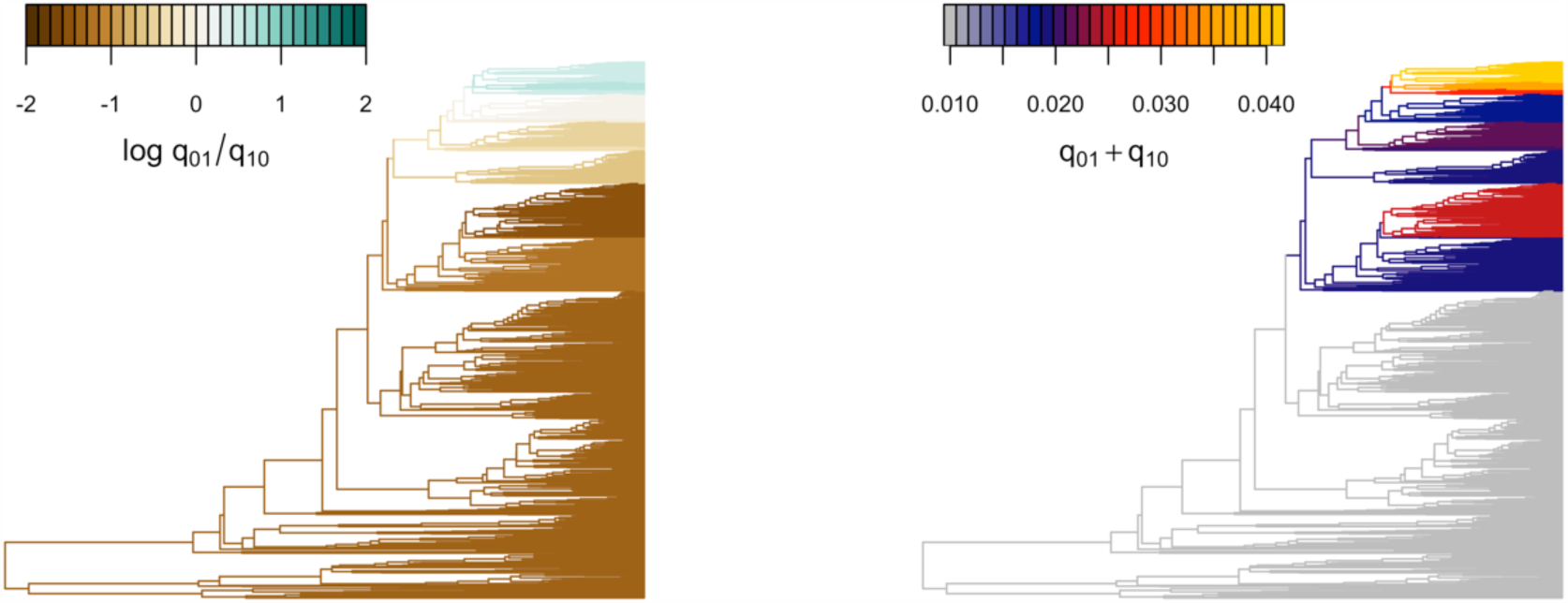
Evolutionary gains and losses of red-black banded coloration show variable tempo and mode across the snake tree of life. Rate matrices were estimated using maximum likelihood conditioned on histories of character evolution inferred with parsimony for clades with at least 10 parsimony-inferred changes. Each clade-specific rate matrix was then compared to the rate-matrix for the whole tree. If the log-likelihood of the data in the subtree formed by a clade was at least 4 log-likelihood units greater under its clade-specific rate matrix than under the tree-wide rate matrix its branches were colored according to the parameters of that matrix. Data from Davis Rabosky et al. (2016).

## Discussion

In this study we show that maximum average likelihood estimates of transition rates for continuous-time Markov chain models of single-character evolution can be improved by using parsimonious ancestral state reconstructions to inform their calculation. We also show how this approach leads to a simple, fast, and effective means of detecting among-clade variation in tempo and mode of character evolution.

To see how combining parsimony and likelihood might be applied to other problems in macroevolutionary inference consider Pagel’s (1994) test for correlated evolution. If we perform this test without using parsimonious ancestor reconstructions a single co-distributed event of character change can lead to a highly significant result (Maddison and FitzJohn 2015; Uyeda et al. 2018). On the other hand, because the data can be explained by a single co-distributed event of character change such a test using a parsimony-informed likelihood estimator would lead to undefined transition rates because (under parsimony) no reconstructed ancestors would have the requisite states (i.e., 01 and 10) needed to compute the transition probabilities. In some respects, the specter of undefined transition rates is an undesirable property of the PIL approach, but the explicit dependence of transition probabilities on independent evolutionary transitions is an important reminder of the need to consider phylogenetic replication when making statements about correlated evolution.

One of the reasons a parsimony-informed approach may be underutilized is that it conditions transition rate estimates on parsimony reconstructions that assume an equally-weighted cost matrix, and parsimony does not provide any criteria for justifying this assumption (Ree and Donoghue 1999). However, the use of simple parsimony makes the approach conservative. If we detect differences in transition rates among states based on reconstructions derived from a cost matrix that assumes there are no differences, we can reasonably conclude that differences exist (Maddison 1994). Furthermore, the alternative, which is to ignore parsimony, conditions transition rate estimates on histories of character evolution for which there is no empirical evidence, or which could be rejected on the grounds of implausibility. For many macroevolutionary datasets we are not interested in those histories as they are unlikely to affect our conclusions, and it is reasonable to ignore them in the computation of the likelihood. In some cases, conditioning rate estimates on the universe of possible histories may actually be misleading. For example, if we attempt to reconstruct the reproductive mode of the most recent common ancestor of living squamates using a Markov model that ignores parsimony it favors a viviparous ancestor at almost two-to-one odds (Pyron and Burbrink 2014). This result is almost certainly incorrect given what we know about the biology of squamate reproductive mode (Shine 2014). The conventional reason given for the model’s failure in this case is the presence of phylogenetic variation in tempo and mode that the model does not account for (King and Lee 2015). While this may be part of the reason, it is worth noting that if we fit the exact same model but condition transition rate estimates on parsimony reconstructions it now favors an oviparous ancestor at almost twenty-to-one odds, a result that is much more plausible but, from a (unconditioned) Markov chain model’s perspective, not as probable.

So far, the discussion has not addressed why mean-squared error in transition rates is lower using parsimony-informed likelihood compared to maximum average likelihood. We suggest this is in part because conditioning rate estimates on parsimony reconstructions helps to reduce the variance in parameter estimates that comes from incorporating the contributions of all possible pathways of evolution. This seems to be most clearly the case for estimates of *τ*, the sum of forward and reverse transition rates (Fig. 2). The parsimony-informed likelihood estimates show a clear negative bias compared to the constrained maximum average likelihood estimates but also less variance, which drives overall mean-squared down. Interestingly, unconstrained maximum average likelihood estimates of *τ* show a strong positive bias and substantially more variance than either of the two other estimators. This positive bias has been noted before (Beaulieu & O’Meara 2016) and is an intrinsic property of the related discrete-time ML estimators studied by Sanderson (1993).

An unexpected result was the increase in mean-squared error we observed among the unconstrained MAV estimates when we examined only the subset of simulations where an unconstrained (asymmetric) model was favored over a constrained (symmetric) model by a likelihood-ratio test. This is cause for concern because it indicates that the rule we use to select a more complicated model over a simpler model also selects for situations that lead to greater estimation errors. Precisely why we observe a difference in performance of the two estimators is not known, but it appears that conditioning estimates on parsimony reconstructions acts to shrink the estimate of *ε* toward 1, which prevents the more extreme biased estimates observed with the MAV estimator.

Finally, we conclude with some remarks on how the results of this study might be affected by a different simulation procedure. Recall that all simulated datasets were generated such that the probability of a net state change from ancestor to descendant was never more probable than retaining the ancestral state. This does not preclude simulations from having a high rate of character evolution, but it does mean that when the rate of character evolution is high the generating model is close to symmetrical. In this case, the likelihood of the model is not influenced by the polarity of state changes and is a decreasing function of the number of character state changes (Tuffley and Steel 1997; Lewis 2001). This helps explain the good performance of parsimony-informed likelihood even when rates of character evolution are high. The possible pathways of character state change that the model assigns a high likelihood closely align with MP pathways. If we did not impose conservative behavior on our simulations it would create a situation where the outcome of a net state change from ancestor to descendant was more likely than the outcome of retaining the ancestral state for at least some, and potentially a large fraction, of the branches in a phylogeny. We do not believe that this is a plausible description of the macroevolutionary process of character evolution. Nonetheless, even in such a situation it is not immediately clear that the performance of the parsimony-informed likelihood approach discussed here would decrease relative to maximum average likelihood because the loss in phylogenetic signal engendered by such a process seems likely to increase the variance of parameter estimates made without the information added by parsimony.

